# Quantification of biases in predictions of protein stability changes upon mutations

**DOI:** 10.1101/308239

**Authors:** F. Pucci, K. Bernaerts, J. M. Kwasigroch, M. Rooman

## Abstract

Bioinformatics tools that predict protein stability changes upon point mutations have made a lot of progress in the last decades and have become accurate and fast enough to make computational mutagenesis experiments feasible, even on a proteome scale. Despite these achievements, they still suffer from important issues that must be solved to allow further improving their performances and utilizing them to deepen our insights into protein folding and stability mechanisms. One of these problems is their bias towards the learning datasets which, being dominated by destabilizing mutations, causes predictions to be better for destabilizing than for stabilizing mutations.

We thoroughly analyzed the biases in the prediction of folding free energy changes upon point mutations (ΔΔ*G*^0^) and proposed some unbiased solutions. We started by constructing a dataset S^sym^ of experimentally measured ΔΔ*G*^0^s with an equal number of stabilizing and destabilizing mutations, by collecting mutations for which the structure of both the wild type and mutant protein is available. On this balanced dataset, we assessed the performances of fifteen widely used *ΔΔ*G*^0^* predictors. After the astonishing observation that almost all these methods are strongly biased towards destabilizing mutations, especially those that use black-box machine learning, we proposed an elegant way to solve the bias issue by imposing physical symmetries under inverse mutations on the model structure, which we implemented in PoPMuSiC^sym^. This new predictor constitutes an efficient trade-off between accuracy and absence of biases. Some final considerations and suggestions for further improvement of the predictors are discussed.

## 1 Introduction

*De novo* protein design is well known to be an important challenge in protein science. Its achievement would have a considerable impact on a wide series of academic and industrial applications that range from drug design in medicinal chemistry to the development of multi-component protein nanomaterials (Zanghellini (2014); Huang *et al*. (2016); Coluzza (2017)). This goal is far from being reached, even though valuable developments have recently been made. Mutational studies with both experimental and computational techniques have thoroughly deepened our understanding of the mechanisms that drive the folding process. In particular, a lot of computational methods have been developed in the last decades to predict in an efficient way how an amino acid substitution impacts protein stability (Dehouck *et al* (2009, 2011); Guerois *et al* (2002); Quan *et al*. (2016); Capriotti et *al* (2005); Pires *et al*. (2014a,b); Pandurangan *et al*. (2017); Laimer *et al*. (2016); Parthiban *et al*. (2006); Kellog *et al*. (2011); Alford *et al*. (2017); Chen *et al*. (2013); Giollo *et al*. (2014); Cheng *et al*. (2006); Masso and Vaisman (2008, 2014)). They allow limiting extensive mutagenesis experiments and thus save time and money.

The most accurate methods among these are structure-based. They use the three-dimensional (3D) structure of the wild type protein as input for predicting how the folding free energy Δ*G*^0^ gets modified upon point mutations (ΔΔ*G*^0^). All these methods are based on a large variety of models that range from pure machine learning algorithms to more biophysics-oriented approaches where the energetic contributions are appropriately combined. Their average performances, measured by the root mean square deviation between experimental and predicted *ΔΔ*G*^0^* values for datasets that contain on the order of two thousand entries, are reported to be between 1.0 and 1.5 kcal/mol (for previous comparisons of the methods’ performances, see Potapov *et al*. (2009) and Khan and Vihinen (2010)). These results are astonishing if one considers that, despite the complexity of the problem, some of the above mentioned tools predict the ΔΔ*G*^0^ of one mutation in less than a minute. This opens the way to perform computational mutagenesis experiments at the proteomic scale with reasonable accuracy.

Unfortunately, these methods suffer from different drawbacks. Like all machine learning approaches, they are prone to overfitting problems (Hawkins (2004); Cawley and Talbot (2010)), and their results therefore tend to be biased toward the training datasets. The analysis and the correction of biases are of primary importance to get more accurate and reliable methods. However, it is a non-trivial task since biases are usually hidden and require careful work on the model structures and on the cleaning of the training datasets.

A known bias in protein stability prediction comes from the fact that the ensemble of experimentally characterized mutations and as a consequence, the training datasets, are dominated by destabilizing mutations. This implies that the predictors tend to be more accurate for destabilizing than for stabilizing mutations, which is a crucial issue given that the latter are the focus of protein design applications. This issue has been reported in a few investigations (Thiltgen and Goldstein (2012); Fariselli *et al*. (2015); Pucci *et al*. (2015)), but there is not yet a common, generally accepted, way to overcome it. Moreover, biases are not limited to this feature but can involve other characteristics such as the kind of protein or the type of wild type and mutant amino acids, since not all substitutions are sufficiently sampled in the training dataset.

In this paper, we go further into this investigation, and assess the performances of different predictors on a new dataset of mutations with experimentally characterized ΔΔ*G*^0^ values and with known 3D structures of both the wild type and mutant proteins. This dataset is by construction balanced with respect to stabilizing and destabilizing mutations. We showed that imposing physical symmetries to the model structures is an efficient and elegant way to solve the bias problem, as already suggested in a preliminary study (Pucci *et al*. (2015)).

## 2 Methods

### 2.1 Folding stability changes upon mutations

Under the assumption that the protein folding process is a reversible, two-state transition - and thus that the protein does not precipitate or aggregate - the thermodynamic stability of a protein can be measured by its folding free energy Δ*G*^0^, *i.e*. the Gibbs free energy difference between the unfolded and folded states:

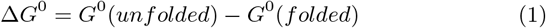

The impact of an amino acid substitution on the protein stability is characterized by the change of Δ*G*^0^ upon mutation

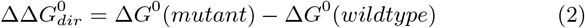

With these conventions, negative values of 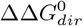 indicate destabilizing mutations while positive 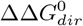 values are associated with stabilizing substitutions. These quantities depend on different thermodynamic and environmental variables such as the temperature and the pH. They are often defined either at room temperature *T_r_* = 25° C or at the melting temperature *T_m_* of the wild type protein. Sometimes, they are not directly measured but derived from Δ*T_m_* measures in differential scanning calorimetry (DSC) experiments, by utilizing the fact that these two quantities are correlated, even though this is only true in a first approximation (see Pucci *et al*. (2016) and Watson *et al*. (2017) for further details). All these dependencies and approximations make the datasets of the experimentally annotated mutations quite noisy, which in turn impacts the accuracy of the predictors that are trained on them.

### 2.2 Assessing predictors through bias evaluation

The change in folding free energy upon mutations is by definition antisymmetric with respect to the exchange between the mutant and wild type residues, assuming that the folding of both the wild type and mutant proteins is a reversible two-state process. This means that the folding free energy 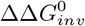 of an inverse mutation, from mutant to wild-type, is equal to minus that of the direct substitution, from wild-type to mutant:

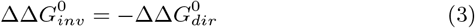

Predictions obtained by computational methods usually do not satisfy this equality, since they are trained on experimental datasets dominated by destabilizing mutations. For example, two of the widely used mutation datasets for model training, S2648 (Dehouck *et al* (2009)) and Q3421 (Quan *et al*. (2016)), exhibit an average ΔΔ*G*^0^ value of −1.01 kcal/mol and −1.13 kcal/mol, respectively. This distortion is learnt by the model and then reproduced in the prediction phase. Note that Eq. (3) cannot be satisfied exactly by the predictors that only consider the wild type and not the mutant structure, but this approximation has usually a small impact when coarse-grained structural representations are used, except in the rare cases where single-site mutations cause large structural rearrangements.

In this paper we constructed a new mutation dataset **S^sym^** which is balanced with respect to stabilizing and destabilizing mutations (see section 2.3), and used it for assessing the performance of fifteen prediction methods (section 2.4) and for quantifying their bias that tends to favor destabilizing mutations. We used the following measures, the former two to estimate the predictors’ accuracy and the latter two the bias:

- *σ_dir_* and *r_dir_* are the root mean square deviation and the linear correlation coefficient between the predicted and experimental ΔΔ*G*^0^ values for the direct mutations in **S^sym^**, from wild type to mutant. Note that these mutations belong to the training dataset of the methods tested, so that the predictions are likely to be overfitted and *σ_dir_* and *r_dir_* to be underestimated and overestimated, respectively.
- *σ_inv_* and *r_inv_* are the root mean square deviation and the linear correlation coefficient between the predicted and experimental ΔΔ*G*^0^ values for the inverse mutations in **S^sym^**, from mutant to wild type. These mutations do not belong to the training datasets and thus constitute an independent test set.
- *r_dir,inv_* is the linear correlation coefficient between the predicted ΔΔ*G*^0^ values of the direct and the inverse mutations. A non-biased prediction that satisfies Eq. (3) has *r_dir,inv_* equal to −1.
- A previously used criterion to estimate the bias is the parameter *δ* defined as (Thiltgen and Goldstein (2012)):

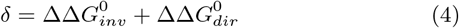

A perfectly non-biased tool should have *δ* = 0 for every mutation. We used here its average value ⟨*δ*⟩ taken over all mutations that belong to **S^sym^**.

## 2.3 Dataset construction

We created a manually curated dataset **S^sym^**, by selecting mutations from the Protherm database (Bava *et al*. (2004)) and checking them on the basis of the original literature. It contains mutations with experimental ΔΔ*G*^0^ values for which the 3D structures of both the wild type and mutant proteins are solved by X-ray crystallography with a resolution of 2.5 Å *atmost*.

Sometimes, different ΔΔ*G*^0^ values are available for the same mutation. We selected the ΔΔ*G*^0^ measured under the environmental conditions closest to the standard conditions (pH=7, *T*=25°C). Note that they are frequently measured at the melting temperature of the wild type protein.

We ended up with a dataset of 684 mutations, half of which are direct mutations inserted in 15 wild type proteins, while the remaining half are inverse mutations inserted in 342 different mutant proteins.

### 2.4 Prediction methods analyzed

We selected the ΔΔ*G*^0^ predictors that are among the most renowned in terms of speed and accuracy. We list them below and briefly explain their characteristics.

1. **PoPMuSiC v2.1** (Dehouck *et al* (2009)): based on standard statistical potentials, combined with sigmoidal weights that depend on the solvent accessibility of the mutated residues.
2. **SDM** (Pandurangan *et al*. (2017)): uses conformationally constrained environment-specific substitution tables to calculate the change in thermodynamic stability between the wild type and the mutant proteins.
3. **CUPSAT** (Parthiban *et al*. (2006)): uses torsion angle potentials and structural environment-specific atom potentials.
4. **Rosetta** (Kellog *et al*. (2011)): generates a 3D structural model of the mutated protein from the wild type structure, and computes the difference in energy between them, with as energy function the sum of a large series of empirical physics-based energy contributions (Coulomb electrostatic, Lennard-Jones atomic interactions, … (Alford *et al*. (2017))).
5. **FoldX v3.0** (Guerois *et al* (2002)): uses a full atomistic description of the protein structure and is based on FOLDEF, an empirical force field developed as a linear combination of different empirical energy terms (van der Waals, solvation, electrostatic, hydrogen bonds …).
6. **I-Mutant v3.0** (Capriotti *et al* (2005)): a tool based on a support vector machine (SVM) that combines protein sequence and structure information.
7. **iSTABLE** (Chen *et al*. (2013)): an integrated predictor, that combines, using an SVM algorithm, sequence information with predictions from different methods (I-Mutant, AUTOMUTE, MUPRO, PoPMuSiC and CUPSAT).
8. **NeEMO** (Giollo *et al*. (2014)): uses an effective representation of proteins based on residue interaction networks (RINs) and combines the extracted information through a neural network.
9. **AUTO-MUTE** (Masso and Vaisman (2008)): uses as main ingredient four-body, knowledge-based, statistical contact potentials that are combined through machine learning tools (random forest and SVM).
10. **STRUM** (Quan et al. (2016)): combines physics- and knowledge-based energy functions derived from protein structure models obtained by I-TASSER (Roy *et al*. (2010)), through gradient boosting regression.
11. **MAESTRO** (Laimer *et al*. (2016)): uses statistical energy functions as main features, and combines them with a multi-agent method that includes a linear regression, an artificial neural network and an SVM.
12. **mCSM** (Pires *et al*. (2014b)): a machine learning method that utilizes graph-based distance patterns between atoms as well as the residue type.
13. **DUET** (Pires *et al*. (2014a)): a consensus prediction method obtained by combining mCSM and SDM using a SVM algorithm.
14. **MUPRO** (Cheng *et al*. (2006)): uses an SVM approach that takes into account sequence information only.

All the tools in this list utilize the 3D structure of the wild type protein as input, except the last one which is based on the protein sequence only. The first five predictors are based on combinations of energy contributions and do not use machine learning, or use machine learning just to identify the parameters of a pre-established model structure. The last nine predictors are true machine learning methods.

Some predictors require as input the pH at which the change in folding free energy is computed (Method 11) or both the pH and the temperature (Methods 5-10), while the others do not ask for the specification of the environmental conditions, assuming standard conditions.

### 2.5 Designing unbiased prediction models

Two approaches can be devised to solve the bias problem and recover predictions that satisfy Eq. (3). One solution is to train the model on a balanced dataset that contains, for each mutation, both the direct and inverse versions, from wild type to mutant and from mutant to wild type. However, this requires knowing the 3D structure of the mutant proteins, which is only available for a subset of mutations: our dataset **S^sym^** contains 684 mutations, whereas the training datasets for which only the wild type structure is requested contain about 3,000 mutations. The datasets can be increased by including mutant structures obtained through comparative modeling, but this introduces noise into the data. Note that this is the only solution in the case of pure machine learning approaches where the model structure is not established *a priori*.

When the prediction model is pre-established and not obtained through a black-box machine learning technique, it is possible to identify the terms in the model structure that are responsible for the symmetry breaking and appropriately correct them. This is exactly what we did in Pucci *et al*. (2015), where the PoPMuSiC^sym^ model, a symmetrized version of PoP-MuSiC v2.1, was presented.

The model structure of the original PoPMuSiC v2.1 is a combination of sixteen contributions:

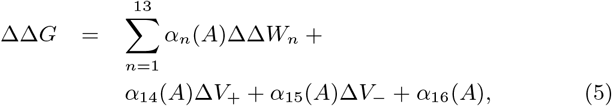

The thirteen terms ΔΔ*W_i_* are changes in folding free energy upon mutation computed using different knowledge-based statistical potentials (see Dehouck *et al* (2009) for details), *α_i_* (i=1.16) are sigmoidal coefficients that depend on the solvent accessibility A of the mutated residues, and Δ*V*± are volume terms defined as:

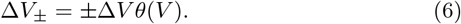

where *θ*(*V*) is the Heaviside function. These two terms represent, respectively, the positive and negative difference in volume between the mutant and wild type amino acids. They provide a description of the impact of the creation of a cavity or the accommodation of stress inside the protein structure. The last term in Eq. (5) is an energy-independent term.

Now, imposing that the model structure satisfies the symmetry relation of Eq. (3) yields the two constraints:

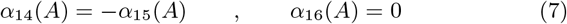

These constraints were introduced into the model structure Eq. (5) and defined a new version of the PoPMuSiC predictor, in which the fourteen remaining *α_i_*(A) parameters were optimized on PoPMuSiC’s original S2648 training dataset. This new version is called PoPMuSiC^sym^ (Pucci *et al*. (2015)).

## 3 Results

We tested fifteen ΔΔ*G*^0^ predictors on a common, balanced, dataset **S^sym^** of 684 single-site mutations, in order to evaluate their performances and, more importantly, their degree of bias with respect to the ΔΔ*G*^0^ symmetry between direct and inverse mutations (Eq. (3)). Table 1 contains the values of the root mean square deviations *σ* and the linear correlation coefficients *r*, reported separately for the direct and inverse mutations. The importance of the bias is evaluated by two parameters, the correlation coefficient *r_dir-inv_* between the direct and inverse mutations and the *δ* parameter defined in Eq. (4).

As clearly seen in Table 1 and Fig. 1, all the tested methods are biased towards the training dataset, except PoPMuSiC^sym^ which has been explicitly designed to avoid this bias. If we focus on direct mutations, the best performing method is MUPRO, a sequence-based machine learning method, with a *σ_dir_* of 0.95 and a *r_dir_* of about 0.8. Remember, however, that all the direct mutations are part of the methods’ training datasets, and these results are thus likely to be affected by overfitting problems. In contrast, the inverse mutations do not belong to the methods’ training sets and can thus be considered as constituting an independent test set. The best performing predictors on the inverse mutations are PoPMuSiC^sym^, MAESTRO, FoldX and PoPMuSiC v2.1.

It is important to note that the black-box machine learning techniques suffer in general more from the bias issue than the other methods that use a more physics-based approach. For example, if one overlooks PoPMuSiC^sym^, the least biased predictor is SDM, which belongs to the physics-based class of predictors, with a correlation coefficient *r_dir-inv_* of about −0.8 and a ⟨*δ*⟩ value of about −0.3.

**Figure 1:**
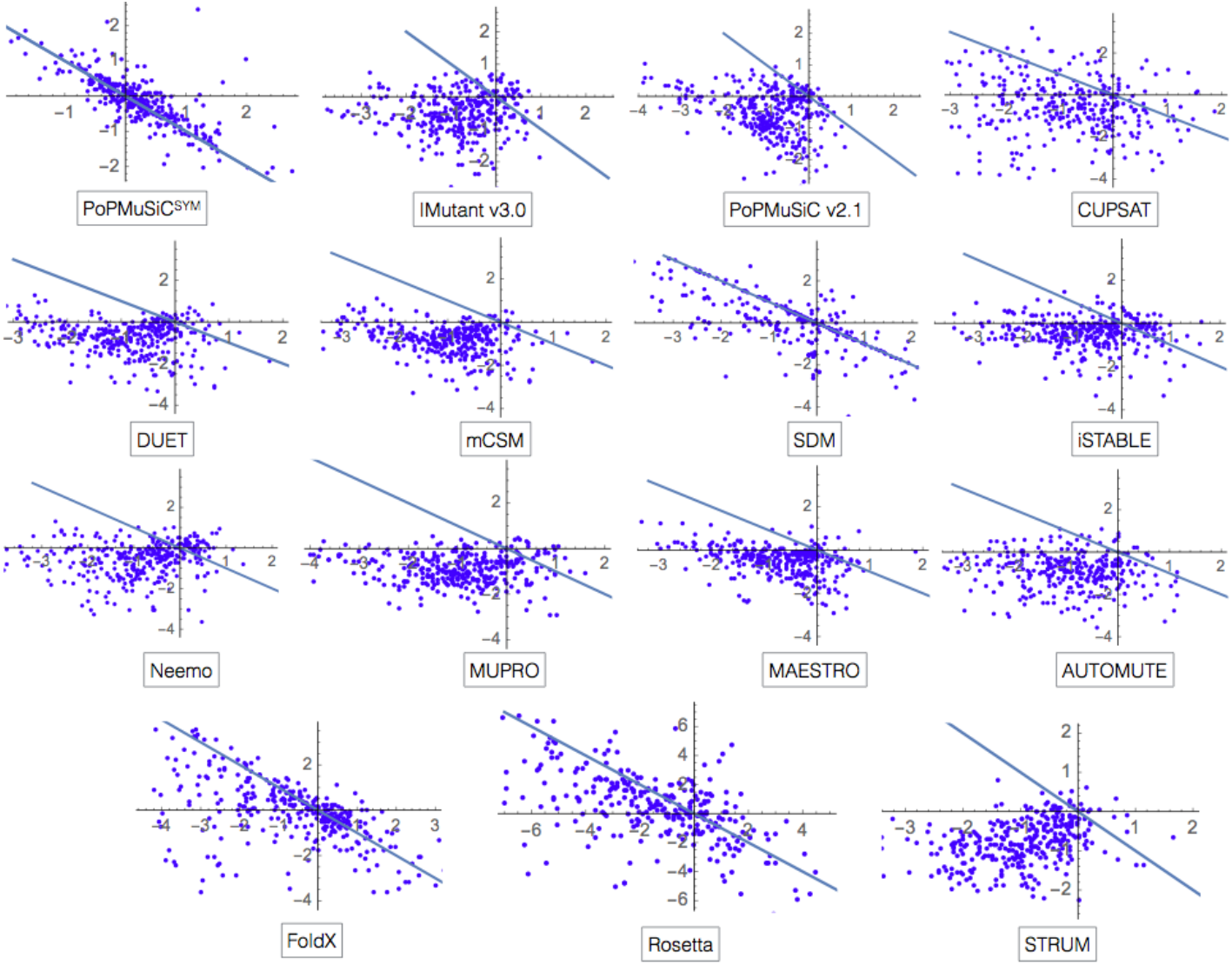
ΔΔ*G*^0^ values (in kcal/mol) of all the mutations in S predicted by the fifteen tools analyzed. The 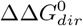 values of the direct mutations (wild type → mutant) are given on the x-axes, and the 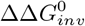 values of the associated inverse mutations (mutant → wild type) are reported on the y-axis. The lines represents the bisectors of the second and fourth quadrants; the perfectly symmetric predictions are on that line.

**Figure 2:**
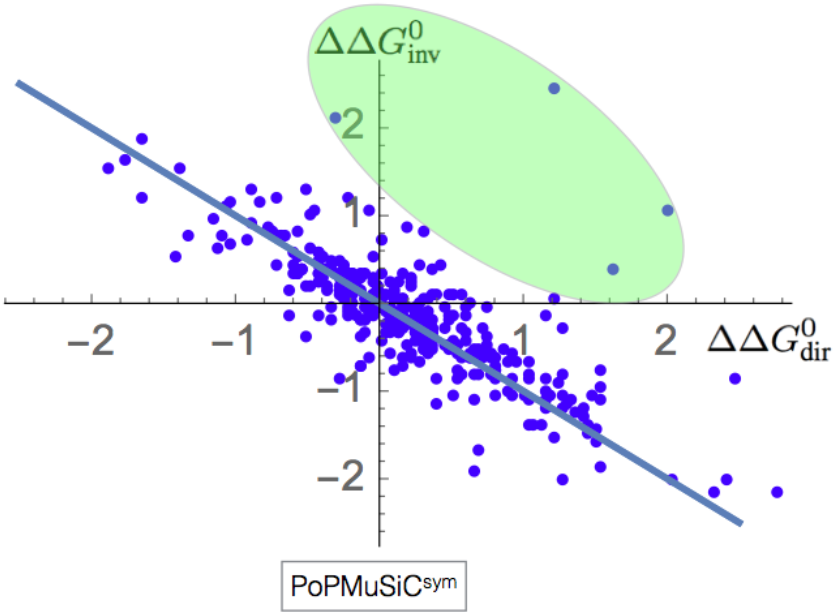
ΔΔ*G*^0^ values (in kcal/mol) of all the mutations in **S** predicted by PoPMuSiC^sym^. The outliers with respect to the symmetric prediction fall in the (green) ellipsoid. They correspond to the four pairs of direct and inverse mutations: (1EY0 K116G; 1KAB G116K), (1EY0 P117A; 1SYG A117P), (1EY0 P117G; 1SYC G117P), (1EY0 P117G; 1SYC G117P), (1EY0 P117T; 1SYE T117P).

However, some physics-based methods are also strongly biased. The point is that such methods can avoid biases only if their model structure is adequately constrained to avoid them. More specifically, the current PoPMuSiC v2.1 version already shows a good performance compared to other predictors, but the implementation of the physical constraints of Eq. (7) in PoPMuSiC^sym^ spectacularly improves *σ_inv_* by more than 25% and yields a zero ⟨*δ*⟩ value.

Note that despite the symmetry constraints there are still some outliers in PoPMuSiC^sym^ with respect to the expected ΔΔ*G*^0^ symmetry between direct and inverse mutations, as shown in Fig. 2. These outliers actually correspond to mutations that cannot be predicted simply from the wild type structure. Indeed, they provoke significant structural rearrangements to avoid steric clashes, empty cavities, or other unfavorable conformations. In these cases, both the wild type and mutant structures should be considered in the ΔΔ*G*^0^ estimation. These issues explain why PoPMuSiC^sym^ does not perfectly satisfy the symmetry relation of Eq. (3) despite its symmetric model structure; the r_dir-inv_ correlation is indeed equal to −0.77 rather than −1.0.

We also analyzed the bias effect separately for core and surface residues. Table 2 reports the results for the best performing methods. In general, the predictions are biased for both surface and core mutations. To correctly interpret these results, we have to remember that mutations in the core have on the average a larger effect on the protein structure and stability. In the **S^sym^** dataset for example, the mean of the absolute values of the ΔΔ*G*^0^s is equal to 1.75 kcal/mol for core mutations and approximatively half (0.96 kcal/mol) for surface mutations. As a consequence, the ⟨*δ*⟩ values of the different methods tend to be worse in the core whereas the *r_dir—inv_* correlations tend to be worse on the surface.

**Table 1:** Bias analysis of all the mutations belonging to the dataset **S^sym^**. The standard deviations *σ*_dir_ and *σ*_inv_ and the values of ⟨*δ*⟩ are in kcal/mol. The methods are ranked according to their performance on the independent test set of inverse mutations, more specifically on the basis of *σ*_inv_.

| Method | $\sigma_{dir}$ | $r_{dir}$ | $\sigma_{inv}$ | $r_{inv}$ | $r_{dir-inv}$ | $\langle\delta\rangle$ |
| --- | --- | --- | --- | --- | --- | --- |
| PoPMuSiC<sup>sym</sup> | 1.58 | 0.48 | **1.62** | **0.48** | **-0.77** | **0.03** |
| MAESTRO | 1.36 | 0.52 | 2.09 | 0.32 | -0.34 | -0.58 |
| FoldX | 1.56 | 0.63 | 2.13 | 0.39 | -0.38 | -0.47 |
| PoPMuSiC v2.1 | 1.21 | 0.63 | 2.18 | 0.25 | -0.29 | -0.71 |
| SDM | 1.74 | 0.51 | 2.28 | 0.32 | -0.75 | -0.32 |
| iSTABLE | 1.10 | 0.72 | 2.28 | -0.08 | -0.05 | -0.60 |
| I-Mutant v3.0 | 1.23 | 0.62 | 2.32 | -0.04 | 0.02 | -0.68 |
| NeEMO | 1.08 | 0.72 | 2.35 | 0.02 | 0.09 | -0.60 |
| DUET | 1.20 | 0.63 | 2.38 | 0.13 | -0.21 | -0.84 |
| mCSM | 1.23 | 0.61 | 2.43 | 0.14 | -0.26 | -0.91 |
| MUPRO | **0.94** | **0.79** | 2.51 | 0.07 | -0.02 | -0.97 |
| STRUM | 1.05 | 0.75 | 2.51 | -0.15 | 0.34 | -0.87 |
| Rosetta | 2.31 | 0.69 | 2.61 | 0.43 | -0.41 | -0.69 |
| AUTOMUTE | 1.07 | 0.73 | 2.61 | -0.01 | -0.06 | -0.99 |
| CUPSAT | 1.71 | 0.39 | 2.88 | 0.05 | -0.54 | -0.72 |

According to our results, the least biased predictors are PoPMuSiC^sym^ and SDM, for both core and surface mutations. But the performance of PoPMuSiC^sym^ is generally better than that of SDM, especially when it is evaluated on the inverse mutation set which does not overlap with the methods’ training sets. The second best performing predictors on the set of inverse mutations is FoldX on core mutations and PoPMuSiC v2.1 on surface mutations.

The bias was also compared between mutations in which an amino acid is replaced by a much larger or a much smaller amino acid, and mutations in which the wild type and mutant amino acids have roughly the same size (Table 3). The volume differences can indeed be another source of bias for some of the prediction methods. Here also, PoPMuSiC^sym^ is the least biased predictor and the best performing on the set of inverse mutations, both for mutations with and without significant size difference. The next least biased predictor is SDM, and the next best performing predictors are MAESTRO and SDM for substitutions with large volume changes, and MAESTRO and FoldX for small volume changes.

**Table 2:** Bias analysis for the 5 best predictors according to the residue localization (core vs surface). The standard deviations *σ_dir_* and *σ_inv_* and the values of ⟨*δ*⟩ are in kcal/mol. The predictors are ranked according to the smallest *σ*_inv_ scores, computed on the set of inverse mutations which constitutes an independent test set, with no overlap with the methods’ training datasets.

| Method | $\sigma_{dir}$ | $r_{dir}$ | $\sigma_{inv}$ | $r_{inv}$ | $r_{dir-inv}$ | $\langle\delta\rangle$ |
| --- | --- | --- | --- | --- | --- | --- |
| **Core Residues** |  |  |  |  |  |  |
| PoPMuSiC<sup>sym</sup> | 1.92 | 0.56 | **1.99** | **0.52** | **-0.89** | **0.03** |
| FoldX | 1.50 | 0.64 | 2.27 | **0.52** | -0.37 | -0.60 |
| SDM | 1.75 | 0.62 | 2.52 | 0.49 | -0.90 | -0.56 |
| MAESTRO | 1.55 | 0.49 | 2.57 | 0.47 | -0.58 | -0.82 |
| PoPMuSiC v2.1 | **1.31** | **0.65** | 2.74 | 0.51 | -0.79 | -1.09 |
| **Surface Residues** |  |  |  |  |  |  |
| PoPMuSiC<sup>sym</sup> | 1.16 | 0.42 | **1.15** | **0.48** | -0.62 | **0.03** |
| PoPMuSiC v2.1 | **1.09** | 0.45 | 1.42 | 0.29 | -0.27 | -0.35 |
| MAESTRO | 1.14 | 0.39 | 1.50 | 0.25 | -0.14 | -0.35 |
| FoldX | 1.61 | **0.60** | 2.00 | 0.18 | -0.39 | -0.35 |
| SDM | 1.72 | 0.18 | 2.02 | 0.16 | **-0.63** | -0.08 |

**Table 3:** Bias analysis for the 5 best predictors according to the difference in volume between wild type and mutant residues. The standard deviations *σ*_dir_ and *σ*_inv_ and the values of ⟨*δ*⟩ are in kcal/mol. The predictors are ranked according to the smallest *σ_inv_* scores, computed on the set of inverse mutations which constitutes an independent test set, with no overlap with the methods’ training datasets.

| Method | $\sigma_{dir}$ | $r_{dir}$ | $\sigma_{inv}$ | $r_{inv}$ | $r_{dir-inv}$ | $\langle\delta\rangle$ |
| --- | --- | --- | --- | --- | --- | --- |
| **Large Volume Changes** |  |  |  |  |  |  |
| PoPMuSiC<sup>sym</sup> | 2.04 | 0.53 | **2.13** | **0.52** | **-0.73** | **0.07** |
| SDM | 1.78 | 0.59 | 2.77 | 0.36 | -0.67 | -0.60 |
| MAESTRO | 1.63 | 0.61 | 2.77 | 0.47 | -0.54 | -0.72 |
| FoldX | 1.89 | 0.60 | 2.90 | 0.41 | -0.27 | -0.84 |
| PoPMuSiC v2.1 | **1.33** | **0.70** | 2.90 | 0.32 | -0.51 | -1.02 |
| **Small Volume Changes** |  |  |  |  |  |  |
| PoPMuSiC<sup>sym</sup> | 1.40 | 0.42 | **1.41** | **0.40** | -0.78 | **0.02** |
| MAESTRO | 1.26 | 0.40 | 1.82 | 0.25 | -0.22 | -0.53 |
| FoldX | 1.44 | **0.60** | 1.83 | 0.36 | -0.46 | -0.35 |
| PoPMuSiC v2.1 | **1.17** | 0.51 | 1.90 | 0.20 | -0.08 | -0.61 |
| SDM | 1.72 | 0.40 | 2.10 | 0.28 | **-0.80** | -0.22 |

## 4 Discussion

In this paper, we thoroughly investigated the ΔΔ*G*^0^ symmetry breaking issue and extensively discussed the fact that computational methods tend to predict the mutations more often as destabilizing than as stabilizing since the training datasets are dominated by destabilizing residue substitutions. Even though this problem was already described in the literature (Thiltgen and Goldstein (2012); Fariselli *et al*. (2015); Pucci *et al*. (2015)), a quantitative measure of the violation of the symmetry between the direct and the inverse substitutions by existing predictors was lacking. This gap has been filled in this paper, in which we quantified and discussed the performance and biases of fifteen of the most efficient available tools. Our results can be summarized as follows:

- All tested methods are biased towards destabilizing mutations. As a proof of this statement, we observed a prediction error on the set of direct mutations (dominated by destabilizing mutations, representing 75% of the dataset entries) which is larger by a factor of about two than the prediction error on the set of inverse mutations (dominated by 75% stabilizing mutations). Indeed, *σ_dir_* is equal to 0.94-1.75 kcal/mol, and *σ_inv_* to 2.09-2.88 kcal/mol. This effect is amplified for the substitutions in the core with respect to surface mutations.
- Predictions that use black-box machine learning techniques tend to be more biased than the others. Indeed, four of the top five prediction tools, PoPMuSiC^sym^, PoPMuSiC v2.1, FoldX and SDM, use biophysics oriented models that combine energy contributions in a coherent way. In contrast, the fifth tool, MAESTRO, uses statistical potentials and other biophysical features combined through several kinds of machine learning methods.
- Imposing biophysical constraints on the model structure (when accessible) is an elegant and simple way to solve completely the bias problem. Indeed, from the analysis of the different folding free energy contributions, it is quite simple to avoid all the terms that violate the symmetry. Relying on symmetry principles in the construction of a model is a common and well known strategy used in physics, which also pays off here, as shown by the spectacular improvement of the S and r_dir-inv_ values of PoPMuSiC^sym^.

Besides the necessity of getting rid of the ΔΔ*G*^0^ symmetry biases, other issues need to be tackled to improve the protein stability prediction methods:

- We would like to draw the attention on the training datasets. Most ΔΔ*G*^0^ predictors use S2648 (Dehouck *et al* (2009)) or Q3421 (Quan *et al*. (2016)) as training sets. These sets are manually curated and based on data coming from the ProTherm database (Bava *et al*. (2004)), which has not been updated since more than five years. As many experimental data have been published since then, especially from deep mutagenesis scanning experiments (Fowler and Fields (2014)), it would be extremely useful to collect them into a new, extended and manually curated database.
- The bias towards destabilizing mutations in the usual learning sets should be taken into account in the evaluation of the methods’ performances. A possibility is to systematically test new methods on S^sym^, the dataset described in this paper that contains both the direct and inverse versions of each mutation and is thus by construction balanced with respect to stabilizing and destabilizing mutations.
- The predictors possibly also suffer from other hidden biases. For example, some types of mutations could be insufficiently sampled in the learning set, with the consequence that the predictor could learn incorrect trends. We would like to stress once more that testing predictors in cross validation is insufficient to correctly evaluate them with respect to the learning dataset biases.
- We would also like to underline the issues related to the addition of more and more features to the predictors. From one side, it allows taking into account the huge complexity of the problem, but from the other side it increases the risk of overfitting and biasing. Moreover, when features are added on top of other features, for example in the case of metapredictors, the performances are difficult to evaluate in genuine cross validation and should be carefully analyzed.

The improvement that the above analyses are expected to bring is crucial in view of addressing even more challenging issues such as the prediction of the changes in folding free energy upon multiple mutations. Indeed, even though it remains costly, the wide screening of single site mutations can be performed experimentally in a reasonable time, via techniques such as deep mutational scanning (Fowler and Fields (2014)). Computational methods capable of predicting only point mutations could thus become less impacting in the protein design field in the near future and the attention should be more focused on the development of predictors that are able to predict the effect of multiple mutations. Such predictions would moreover be more likely to fulfill the requirements of improving protein stability in biotechnological applications, which are frequently impossible to satisfy by single point mutations only, but require combinations of mutations to achieve, for example, high energetic stabilization while maintaining the solubility and activity of the protein unaltered.

## Supporting Material

- Dataset **S^sym^**

